# A Nascent Peptide Code for Translational Control of mRNA Stability in Human Cells

**DOI:** 10.1101/2021.12.01.470782

**Authors:** Phillip C. Burke, Heungwon Park, Arvind Rasi Subramaniam

**Affiliations:** Basic Sciences Division and Computational Biology Section of the Public Health Sciences Division, Fred Hutchinson Cancer Research Center, Seattle, WA 98109, USA

## Abstract

Stability of eukaryotic mRNAs is associated with their codon, amino acid, and GC content. Yet, coding sequence motifs that predictably alter mRNA stability in human cells remain poorly defined. Here, we develop a massively parallel assay to measure mRNA effects of thousands of synthetic and endogenous coding sequence motifs in human cells. We identify several families of simple dipeptide repeats whose translation triggers acute mRNA instability. Rather than individual amino acids, specific combinations of bulky and positively charged amino acids are critical for the destabilizing effects of dipeptide repeats. Remarkably, dipeptide sequences that form extended β strands *in silico* and *in vitro* drive ribosome stalling and mRNA instability *in vivo*. The resulting nascent peptide code underlies ribosome stalling and mRNA-destabilizing effects of hundreds of endogenous peptide sequences in the human proteome. Our work reveals an intrinsic role for the ribosome as a selectivity filter against the synthesis of bulky and aggregation-prone peptides.

## Introduction

Protein expression is determined by a balance between the translation rate and stability of mRNAs. In human cells, mRNA stability is often regulated by sequence motifs in the 3′ untranslated region such as microRNA-binding sites and AU-rich elements (*1*). Additionally, the protein coding region has been recently recognized as a critical determinant of eukaryotic mRNA stability (*2, 3*). The role of the coding sequence in mRNA stability is best understood in the budding yeast *S. cerevisiae* where poorly translated codons and nascent peptide motifs with positively charged residues can destabilize mRNAs (*4*–*7*). Poorly translated codons have also been implicated in regulation of mRNA stability in several other organisms (*8*–*11*).

Coding sequence features regulating mRNA stability in human cells are less clear. Several recent studies examined the coding sequence determinants of endogenous mRNA stability in human cells and arrived at differing conclusions. Two studies implicated synonymous codon choice as the primary determinant of mRNA stability in human cells (*12, 13*). Another found GC and GC3 (wobble base GC) content as major factors regulating mRNA stability (*14*). A fourth study identified amino acid content to be an important contributor (*15*). Extended amino acid motifs and G-quadruplexes in coding regions have also been implicated as triggers of specific mammalian mRNA decay pathways (*16, 17*). The associations reported in these studies relied on endogenous human coding sequences. Since human mRNAs differ from each other in codon, amino acid, and GC content as well as in their length and the presence of specific sequence motifs, it is challenging to identify the contribution of each factor to mRNA stability. Further, reporters used in the above studies for validation differ extensively in their nucleotide or amino acid content, which complicates their interpretation.

Here, we developed a massively parallel assay to measure the mRNA effects of thousands of coding sequence motifs in human cells. We designed our assay with the initial goal of systematically delineating the individual contribution of mRNA features implicated in previous studies. Instead, we unexpectedly uncovered a potent role for the sequence and structure of the nascent peptide in regulating mRNA stability and ribosome elongation rate. The resulting nascent peptide code regulates the ribosome stalling and mRNA destabilizing effects of hundreds of endogenous peptide sequences from the human proteome. Our results point to an unappreciated role for the ribosome as a selectivity filter against the synthesis of bulky and aggregation-prone peptide sequences.

## Results

### A massively parallel assay for mRNA levels in human cells

We reasoned that coding sequence motifs that alter mRNA stability should be identifiable through their effects on steady state mRNA levels. To study the effect of coding sequence motifs on mRNA levels in an unbiased manner, we designed a library of 4,096 oligonucleotides made of all possible codon pairs (Fig. 1A). We repeated each codon pair as a tandem 8× repeat with the rationale that their effects will be amplified and readily measurable. We cloned the oligonucleotide library as a pool into a dual fluorescence reporter vector separated by 2A linkers – a design widely used for studying ribosome stalling motifs in human cells (*18*–*23*). We added multiple random 24nt barcodes without stop codons 3′ of each oligonucleotide insert and linked the barcode sequences to the corresponding insert by high-throughput sequencing. Most studies of coding sequence motifs use transient transfection or lentiviral integration of reporters, which makes measurement of steady state effects on mRNA levels across a large pool difficult. To avoid this, we stably integrated the reporter pool at the *AAVS1* locus of HEK293T cells using CRISPR Cas9-mediated homologous recombination. We extracted mRNA and genomic DNA from the pooled cells and counted each barcode by high-throughput sequencing. Normalization of the total barcode count in the mRNA by the corresponding count in the genomic DNA for each of the 4,096 inserts provides a relative measure of the steady-state mRNA level of that insert.

**Figure 1.**
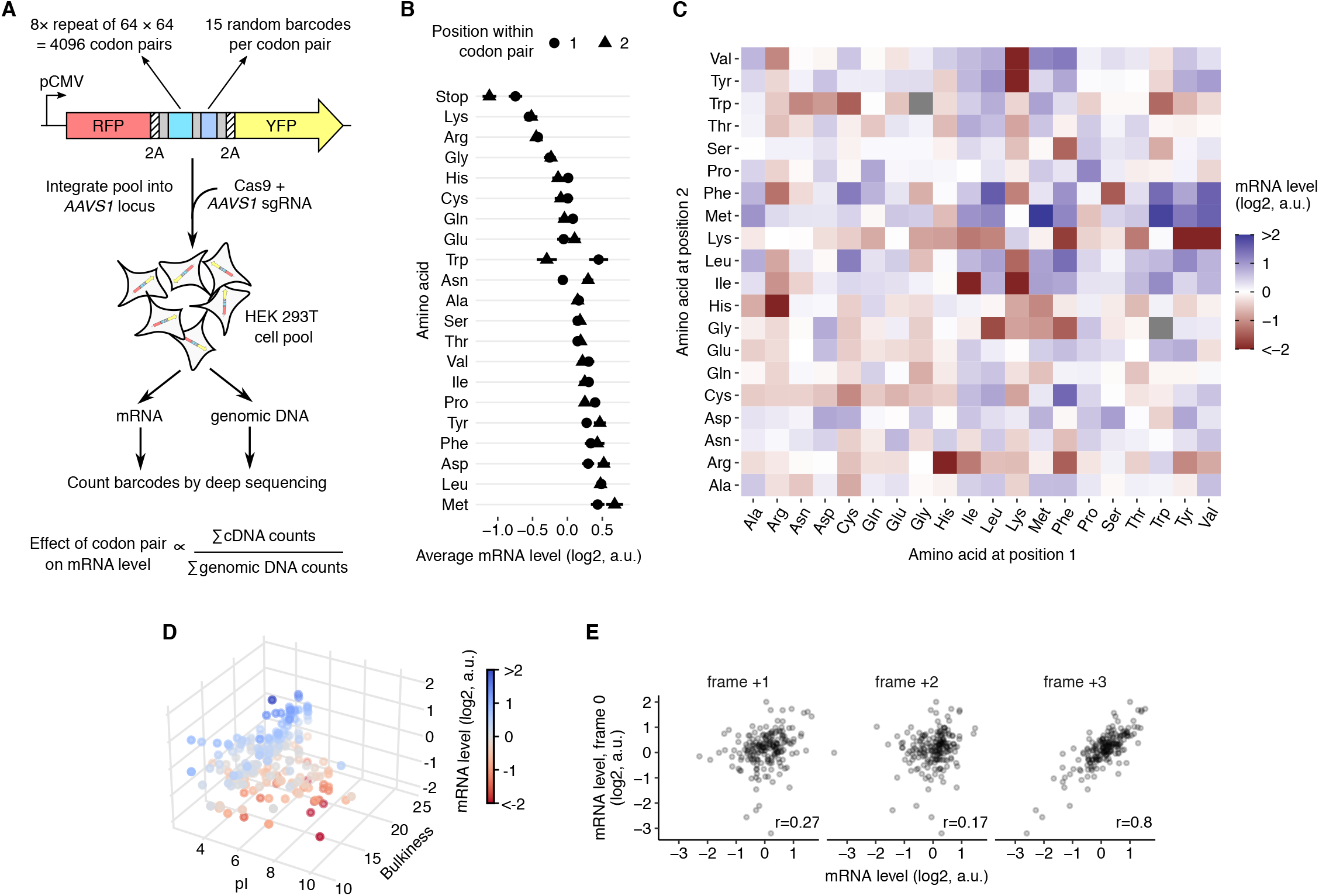
Dipeptide repeats trigger mRNA instability. **(A)** Schematic of massively parallel assay for measuring reporter mRNA levels. 8× repeats of all 4096 codon pairs are synthesized as pooled oligonucleotides, linked in-frame to 24nt random barcodes, and cloned between RFP and YFP reporters with intervening 2A sequences. Each insert has a median of 15 random barcodes without in-frame stop codons. Reporter cassettes are integrated as a pool at the *AAVS1* locus in HEK293T cells by Cas9-mediated homologous recombination and constitutively expressed off the CMV promoter. Steady state mRNA level of each insert is determined by sequencing corresponding barcodes in the cDNA and the genomic DNA and normalizing the summed cDNA read counts by the genomic DNA read counts. **(B)** mRNA level of reporters with codons encoding one of the twenty amino acids or a stop codon in position 1 (circles) or position 2 (triangles) of the 8× codon pair insert shown in A. **(C)** mRNA level of reporters encoding 400 different dipeptide repeats. Amino acids encoded by the first or second position in the dipeptide are shown along the horizontal or vertical axis respectively. Two dipeptide repeats with missing values are shown in grey. **(D)** mRNA level of dipeptide repeat-encoding reporters plotted as a function of the average isoelectric point (pI) and the bulkiness (*32*) of the two amino acids in the dipeptide. **(E)** mRNA level of reporters encoding 190 different dipeptide repeats (excluding reversed repeats) in the correct reading frame (frame 0, vertical axis) or in reading frames shifited by +1, +2, or +3 nucleotides (horizontal axes). *r* is the Pearson correlation coefficient between frame 0 and the frame-shifited mRNA levels. mRNA levels in B–E are in arbitrary units (a.u.) and are normalized to the median value across all dipeptide repeats. Error bars in B represent standard errors of measurement over barcoded replicates calculated by bootstrap. Most error bars in B are smaller than data markers. ^**^ and ^*^ indicate one-sided t-test P-values of <0.01 and <0.05 respectively for decrease in mRNA levels at indicated time points of the destabilizing dipeptide repeats with respect to their +2 frameshifit controls.

We examined whether our assay captured the effects of known mRNA-destabilizing motifs. Stop codons in either the first or second position of the codon pair repeat decrease mRNA levels (Fig. 1B), consistent with their mRNA destabilizing effect due to nonsense-mediated decay (*24*–*26*). However, the destabilizing effects of codon pairs are only weakly correlated with known effects of GC content, synonymous codon usage, or amino acid content on mRNA stability (Fig. S1A-C) (*12*–*15*). Among the twenty amino acids, the positively charged amino acids lysine and arginine cause the largest average decreases in mRNA levels (Fig. 1B). This is in line with the known association between positively charged residues in the nascent peptide and slow elongation (*27* –*31*).

### Specific dipeptide repeats trigger mRNA instability

We wondered whether the mild average effects of amino acids on mRNA levels (Fig. 1B) belie larger effects driven by specific amino acid combinations. Therefore, we assessed the effect of each pairwise amino acid combination on mRNA level (Fig. 1C). While lysine and arginine are destabilizing on average (Fig. 1B), unexpectedly, these amino acids have mild or no destabilizing effect on mRNA levels on their own (Fig. 1C: Lys-Lys, Arg-Arg, Arg-Lys). Rather, the destabilizing effects of lysine and arginine are primarily driven by co-occurrence with bulky amino acids (*32*) (ratio of side chain volume to length > 18Å^2^) such as valine, isoleucine, leucine, phenylalanine, and tyrosine (Fig. 1C). Likewise, most bulky amino acids are highly destabilizing in combination with lysine and arginine, but not on their own (Fig. 1C). Further, a few dipeptides that contain certain positively charged amino acids (Arg-His) or bulky amino acids (Phe-Ser) also have a strong mRNA destabilizing effect (Fig. 1C). The combinatorial effect of positively charged and bulky amino acids on mRNA level is captured by a linear statistical model (Fig. 1D): Isoelectric point (*32*) (pI, a measure of positive charge) and bulkiness (*32*) of amino acids are positive correlates of mRNA level, while an interaction term between these two physical properties is a negative correlate of mRNA level [mRNA = (0.31 × pI) + (0.20 × bulkiness) – (0.03 × pI × bulkiness), Adjusted R^2^ = 0.25]. By contrast, ignoring the interaction between pI and bulkiness results in negative or no correlation of these properties with mRNA level (mRNA = – 0.18 × pI, Adjusted R^2^ = 0.21), which is in line with Fig. 1B. The destabilizing effects of dipeptide repeats in the translated +0 frame strongly correlates with the codon-matched +3 frame, but only weakly with the codon-mismatched +1 and +2 frames (Fig. 1E). The high correlation between the +0 and +3 frames is also seen from the diagonal symmetry of Fig. 1C and arises from similarity of the encoded peptides (for eg. (XY)_8_ and (YX)_8_ are identical except at their termini). These frame correlations are consistent with the mRNA destabilizing effects arising at the translational level as opposed to transcriptional or RNA processing differences. Together, our results show that translation of bulky and positively charged amino acids is critical for their negative effect on mRNA level.

### Nascent peptide primary sequence modulates mRNA stability

Several observations suggest that translation of specific dipeptide repeats is a general trigger of mRNA instability in human cells. First, mRNA levels of destabilizing repeats are lower than their frameshifted controls in multiple human cell lines (HEK293T, HeLa, HCT116, and K562; Fig. 2A), pointing to the generality of the observed mRNA effects. Second, upon actinomycin D treatment to inhibit transcription, mRNA levels of reporters with destabilizing dipeptides decay faster than their frameshifted controls (Fig. 2B). This confirms that the decrease in steady-state mRNA levels caused by dipeptide repeats arises from reduction in mRNA stability. Third, to decipher the effect of dipeptide repetition on mRNA instability, we systematically varied the number of several destabilizing dipeptides identified in our initial assay (Fig. 2C). As the number of dipeptide repeats increases from 1 to 8, each dipeptide becomes destabilizing at a distinct repeat number between 4 and 7 (Fig. 2C). Fourth, we altered the periodicity of dipeptide repeats by intermixing dipeptides with their reversed counterparts such that the overall amino acid composition remains unchanged (Fig. 2D). Even minor perturbations of RH repeats completely abrogate their mRNA destabilizing effect (Fig. 2D, upper panel). VK repeats become gradually less destabilizing as their periodicity is decreased, while SF repeats show an intermediate trend (Fig. 2D, middle and lower panels). This last experiment reveals that the primary sequences of dipeptide repeats encode critical regulatory information beyond the identity of the amino acid pairs forming the repeats.

**Figure 2.**
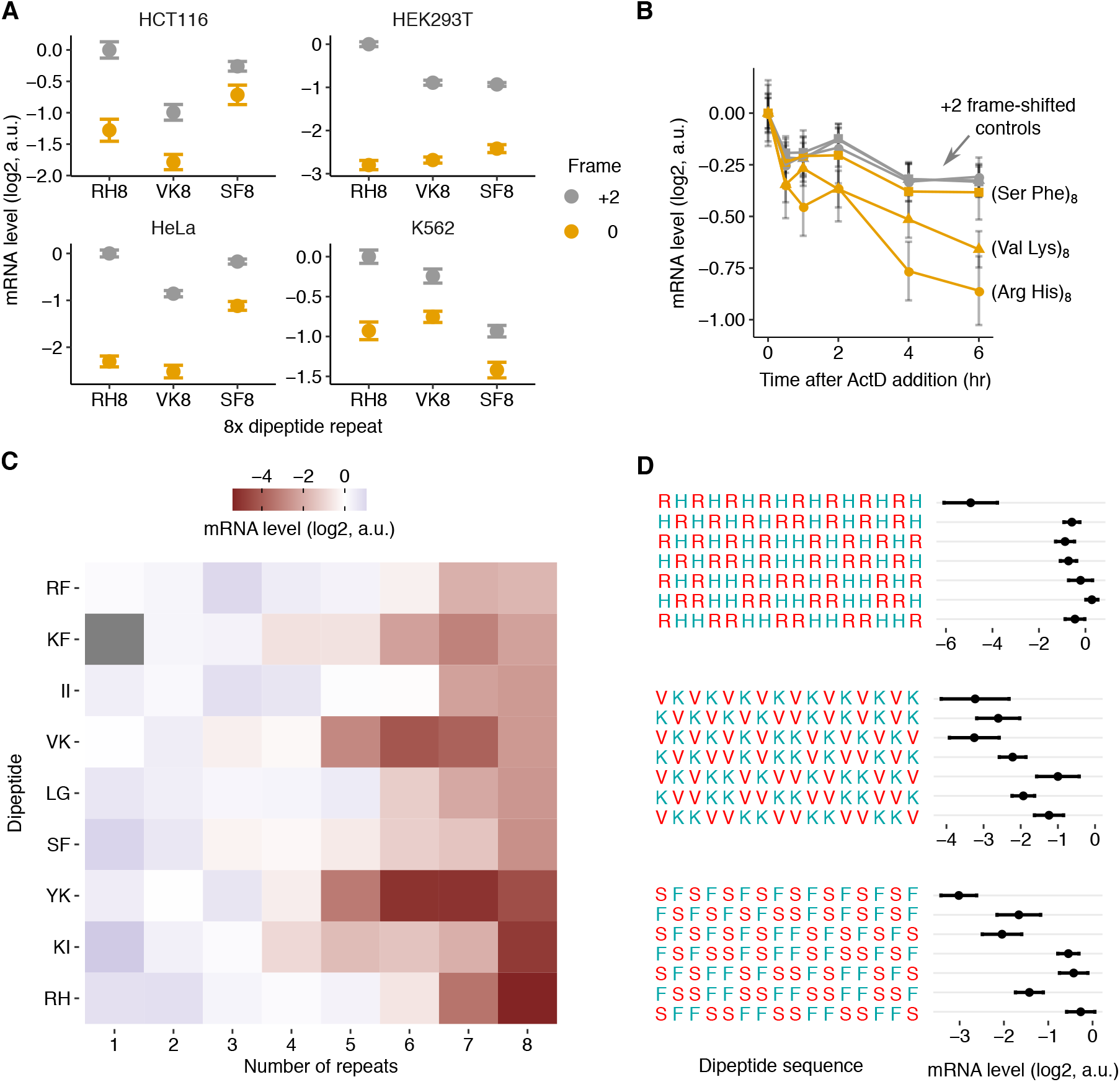
Nascent peptide primary sequence modulates mRNA stability in human cells. **(A)** mRNA levels of reporters with dipeptide repeats (orange): (Arg His)_8_, (Val Lys)_8_, (Ser Phe)_8_, or the three +2 frameshift controls (grey): (Pro Ser)_8_, (Gln Ser)_8_, (Phe Gln)_8_ in 4 different cell lines: HCT116, HEK293T, HeLa, and K562. **(B)** mRNA stability of reporters from **A** in HEK293T cells. Reporter mRNA levels are measured at indicated time points after Actinomycin D-induced transcriptional shut off. Most points for the three frameshift controls overlap with each other. **(C)** mRNA levels of dipeptide-encoding reporters with different dipeptide repeat length. Missing value is shown in grey. **(D)** mRNA levels of dipeptide-encoding reporters with different dipeptide repeat periodicity. Periodicity of dipeptides is systematically reduced by interspersing 1, 2, or 4 tandem repeats of each dipeptide with an equal number of its sequence-reversed counterpart. mRNA levels are measured using the pooled sequencing assay in Fig.1A and normalized by the median value across all inserts in the pool. Amino acids in A, C, and D are labeled by their one-letter codes. Error bars in A, B, and D represent standard errors of measurement over barcoded replicates calculated by bootstrap.

### Secondary structure of dipeptide repeats mediates mRNA stability effects

Since dipeptide sequences are known to form distinct secondary structures based on their periodicity (*33, 34*), we asked whether mRNA-destabilizing dipeptide repeats adopt specific secondary structures. Using a deep neural network model for secondary structure prediction (*35*), we assigned dipeptide repeats to either α helices or β strands if their respective prediction probabilities are greater than 0.5. We find that dipeptide repeats predicted to form β strands have a significantly lower mRNA level on average than those predicted to form α helices (Fig. 3A, P < 0.001, two-sided Mann-Whitney test). Further, many dipeptide repeats that strongly destabilize mRNAs *in vivo* are also computationally predicted to form β strands with a high probability (Fig. 3B). Notably, among dipeptides containing the positively charged amino acids lysine or arginine, the measured propensity of the second amino acid to occur in a β strand (*36*) (‘Chou-Fasman propensity’) is highly correlated with mRNA instability (Fig. 3C, left; Fig. S2A). This correlation is not observed with α helix propensities of the same amino acids (Fig. 3C, right; Fig. S2A). mRNA levels of dipeptide repeats containing the negatively charged glutamate, which are also predicted to form β strands with high probability when combined with bulky amino acids (Fig. S2B), do not show significant correlation with β strand or α helix propensities (Fig. S2C). Thus, a combination of bulky and positively charged amino acids in the primary sequence and β strand in the secondary structure are strong and significant predictors of the mRNA-destabilizing effects of dipeptide repeats [mRNA = (0.30 × pI) + (0.23 × bulkiness) – (0.03 × pI × bulkiness) – (0.52 × β-strand-propensity), Adjusted R^2^ = 0.27].

**Figure 3.**
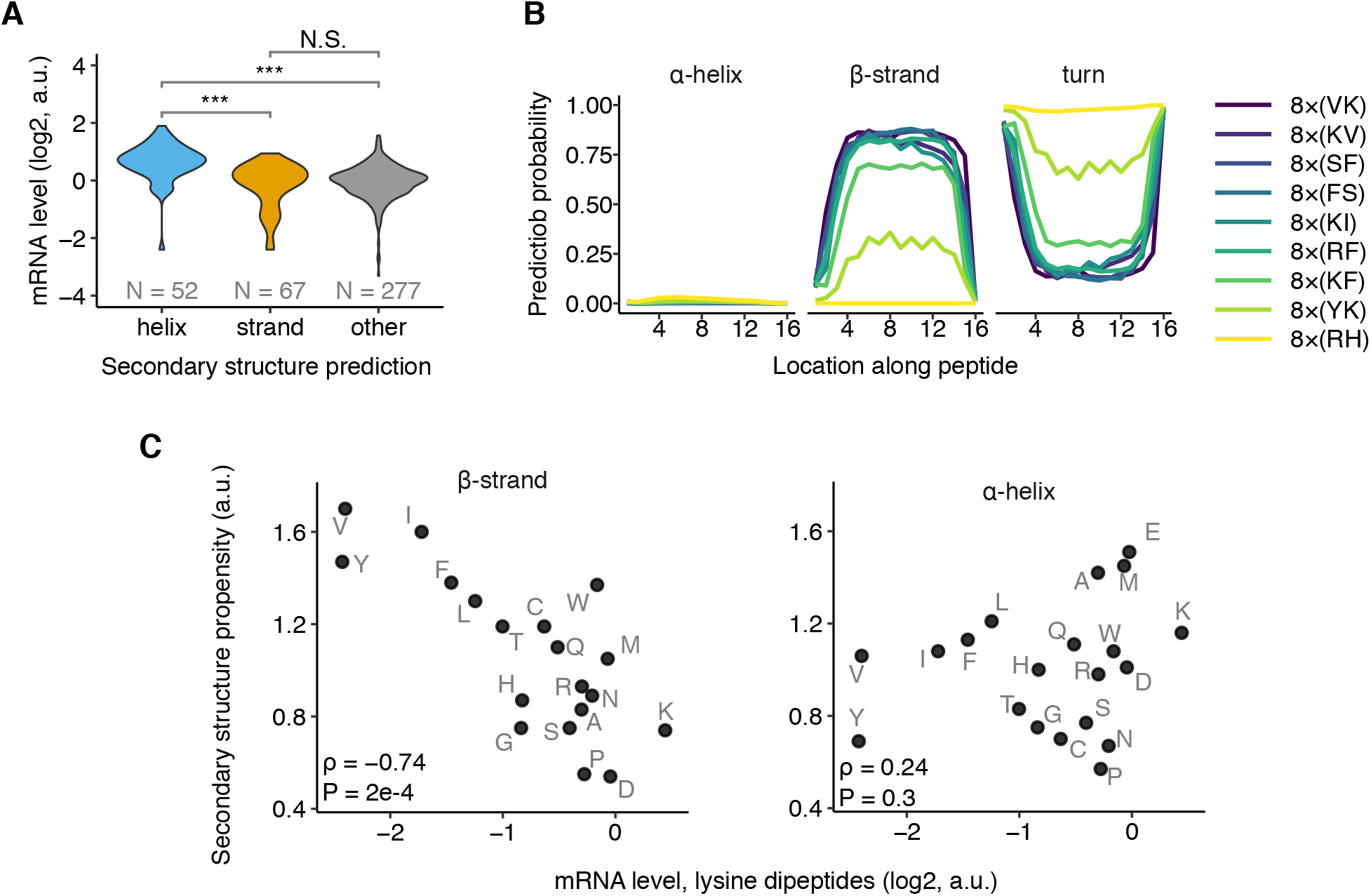
Secondary structure of dipeptide repeats mediates mRNA instability effects. **(A)** Distribution of mRNA levels of dipeptide repeat-encoding reporters from Fig. 1C partitioned by predicted protein secondary structure. Per residue probability of secondary structure formation are predicted using S4PRED (*35*). Inserts with >50% average prediction probability of forming α helix or β strand are classified as such, or else grouped as ‘other.’ *N* is the number of dipeptide repeats predicted to be in each category. ^***^: P < 0.001, N.S.: not significant (two-sided Mann-Whitney U test). **(B)** Computationally predicted secondary structure probability along 16 amino acid-long peptide sequences encoded by destabilizing dipeptides. Secondary structure probabilities are predicted using S4PRED. **(C)** mRNA levels of dipeptide repeat-encoding reporters with lysines in one position and one of twenty amino acids in the other position of the repeat (labeled in grey) shown on horizontal axes. Propensity (*36*) of the second amino acid to occur in a β strand (left) or an α helix (right) shown on vertical axes. ρ is the Spearman correlation coefficient between the two axes with the indicated *P* value.

### Extended β strands drive ribosome stalling and mRNA instability

To test the causal role of β strands in nascent peptide-mediated translational control, we combined the mRNA-destabilizing dipeptides VK, KV, SF, and FS into 16 amino acid-long peptides. Even though the four constituent dipeptides are strongly predicted to form β strands on their own (Fig. 3A), their combinations can form either β strands or α helices with high probability (Fig. 4A). Importantly, all combinations are encoded by the same set of four amino acids to control for amino acid composition. We commercially synthesized two 16 amino acid peptides and used circular dichroism to confirm their secondary structure *in vitro* (Fig. 4B, left panel). As predicted (Fig. 4A), 4×SVKF primarily forms β strands in aqueous solution, while 4×SKVF forms α helices in the presence of trifluoroethanol (TFE) as a co-solvent (*37* –*39*) (Fig. 4B, right panel). We then measured the transit time of ribosomes on mRNAs encoding 16 amino acid inserts preceding a Nanoluciferase reporter in a rabbit reticulocyte lysate (RRL) *in vitro* translation system (Fig. 4C, left). The β strand-forming 4×SVKF and 4×VKFS inserts slow ribosome elongation relative to the α helix-forming 4xSKVF and 4xKVFS inserts (Fig. 4C, middle), with a 200 s difference in *in vitro* transit time (Fig. 4C, right). Strikingly, all β strand peptides decrease mRNA levels over 8-fold relative to α helix controls when tested *in vivo* using our reporter assay (Fig. 4D). We observe similar destabilizing effects due to β strand formation in HeLa, HCT116, and K562 cells (Fig. S3). We conclude that the propensity of nascent peptides to form β strands is a critical determinant of dipeptide-mediated ribosome stalling and mRNA instability in human cells.

**Figure 4.**
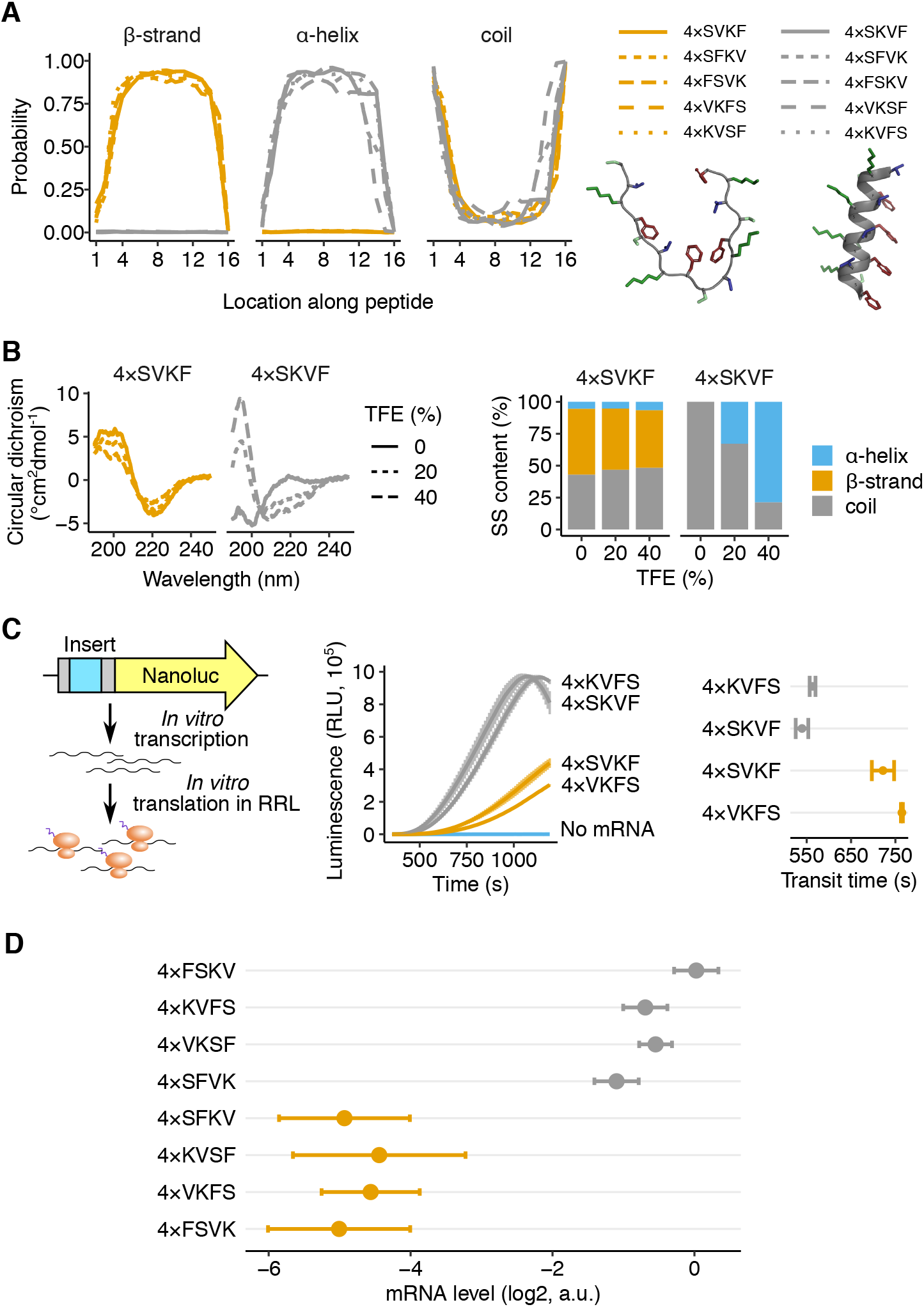
Extended β strands drive ribosome stalling and mRNA instability. **(A)** Computationally predicted secondary structure probability along 16 amino acid-long peptide sequences encoded by alternating VK or KV dipeptides with SF or FS dipeptides identified as destabilizing in Fig. 1. Secondary structure probabilities are predicted using S4PRED. The 10 different peptide sequences are 4× repeats of the dipeptide combinations shown in the legend (for example, 4×SVKF: SVKF SVKF SVKF SVKF). Predicted β strand and α helix structures of 4×SVKF and 4×SKVF respectively using PEP-FOLD3 (*92*) are shown below the legends. The peptide backbones are in grey and the side chains of amino acids are colored. **(B)** Measured circular dichroism spectra of *in vitro* synthesized 4×SVKF or 4×SKVF peptides (left). Measurements are performed with 0, 20, 40% of Trifluoroethanol (TFE) as co-solvent in 10 mM sodium phosphate buffer (pH = 7.5). Relative content of different secondary structures is estimated by linear deconvolution of the measured spectra from a pre-computed basis set using SESCA (*93*). **(C)** *In vitro* measurements of ribosome transit time on mRNAs encoding β strand-or α helix-forming peptides followed by Nanoluciferase (left panel). Luminescence is measured as a function of time after addition of *in vitro* transcribed mRNAs to rabbit reticulocyte lysate (RRL) at t=0*s* (middle panel). Standard error of measurement across three technical replicates is shown as a shaded area on either side of the mean. Ribosome transit times (right panel) are estimated by measuring the X-intercept of the linear portion of the raw luminescence signal in the middle panel. **(D)** *In vivo* mRNA levels of reporters encoding one of eight different dipeptide combinations. mRNA levels are measured using the reporter constructs and pooled sequencing assay in Fig. 1A. Error bars represent standard errors of measurement over barcoded replicates calculated by bootstrap.

### Dipeptide motifs in the human genome trigger ribosome stalling and mRNA instability

We sought to identify endogenous sequences in the human genome that regulate mRNA translation based on the dipeptide code identified above. To do this, we scanned all annotated human protein coding sequences for destabilizing dipeptide combinations of bulky and positively charged amino acids (Fig. 5A). Using a heuristic peptide score (Fig. 5A, top), we identified the 16 amino acid long peptide within each coding sequence that has the maximum density of destabilizing dipeptides. mRNA regions encoding these dipeptide motifs exhibit increased ribosome occupancy relative to control motifs from the same mRNAs in HEK293T cells (*23*) (Fig. 5B). A similar increase in ribosome occupancy is also seen in other human and mouse cell lines (*40, 41*) (Fig. S4A,B), while total RNA density does not show such an increase (Fig. S4A,B). The peak in P-site density at about 25nt from the center of the motif (Fig. 5B) suggests that the 16 amino acid peptides are almost fully synthesized before they stall the ribosome.

**Figure 5.**
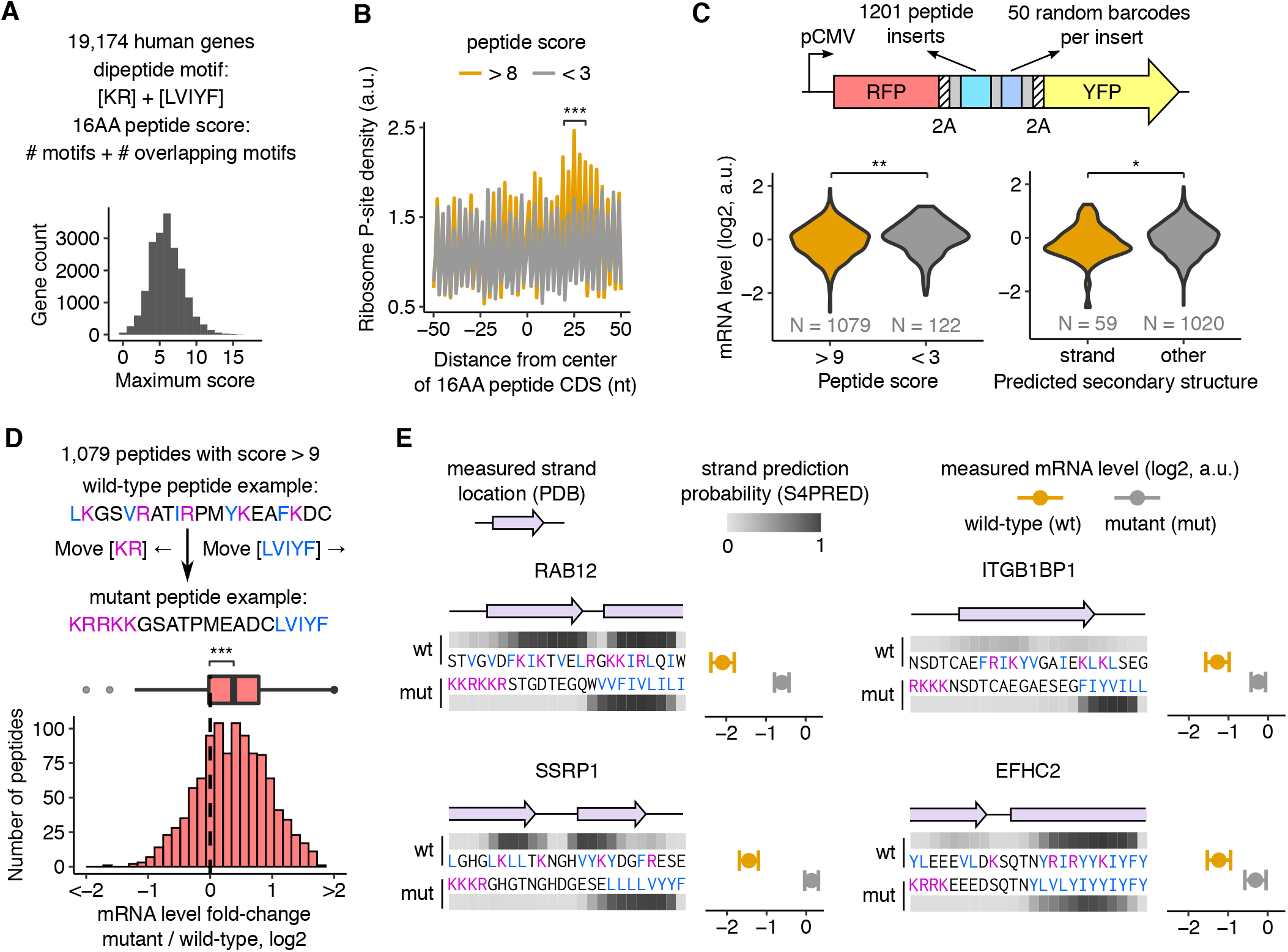
Dipeptide motifs in the human genome trigger ribosome stalling and mRNA instability. **(A)** Scoring methodology for dipeptide motifs in human CDS. Destabilizing dipeptides formed by lysine (K) or arginine (R) with an adjacent leucine (L), valine (V), isoleucine (I), tyrosine (Y), or phenylalanine (F) are given a score of 1. If two such dipeptides overlap, an additional score of 1 is given. The 16 amino acid peptide window with the maximum score is identified in each CDS, and the distribution of these peptide scores across all genes is shown in the lower panel. **(B)** Measured ribosome P-site density around destabilizing dipeptide motifs in HEK293T cells (*23*). The first nucleotide of the 9th codon encoding the 16AA peptide motif is designated as 0nt. ^***^: P = 1e-5 (two-sided Mann-Whitney U test) for increase in ribosome P-site density in +20 to +30nt around the destabilizing peptide motif. Comparison is between motifs with peptide score > 8 and randomly chosen control motifs in the same set of genes with peptide score < 3. 485 motifs in genes with an average ribosome P-site density greater than 1 per codon are included in this metagene analysis. Metagene ribosome P-site densities are averaged across all destabilizing or control motifs after normalization by the mean density for each gene. **(C)** mRNA levels of destabilizing (peptide score > 9) and control motifs (peptide score < 3) are measured by pooled cloning (1,201 total inserts) into a reporter construct followed by deep sequencing as in Fig. 1A. Left panel: Distribution of measured mRNA levels of destabilizing dipeptide motifs compared to control motifs. Right panel: Distribution of measured mRNA levels of destabilizing dipeptide motifs in the left panel partitioned by predicted secondary structure. 59 motifs with an average β strand prediction probability > 0.5 using S4PRED are classified as β strands. **(D)** Increase in measured mRNA levels upon reordering amino acids in 1,079 endogenous destabilizing dipeptide motifs from C (median Δlog2 mRNA = 0.38). All codons encoding K or R are moved to the 5′ end of the mutated motif and codons encoding L, V, I, Y, or F are moved to the 3′ end. mRNA levels of motifs are measured using the pooled reporter assay in C. ^***^: P < 0.001, ^**^: P < 0.01, ^*^: P < 0.05 using two-sided Mann-Whitney U test in C and D. **(E)** Examples of endogenous destabilizing motifs with known β-stranded secondary structure. The measured secondary structure of each wild-type motif from PDB is shown as a purple ribbon diagram. Prediction probability for β strands using S4PRED is shown as a grayscale heatmap for wild-type and mutant motifs. Measured mRNA levels of wild-type (orange) and mutant (grey) motifs are shown on the right within each panel. mRNA levels of wild-type and mutant motifs are measured using the pooled reporter assay in C. Error bars represent standard errors of measurement over barcoded replicates calculated by bootstrap.

To test whether the endogenous motifs identified above can destabilize mRNA, we cloned 1,201 such motifs into our reporter and measured their mRNA levels by high throughput sequencing (Fig. 5C, top). Motifs with high destabilizing dipeptide content result in lower mRNA levels than control motifs (P < 0.01, Fig. 5C, left panel). Among destabilizing motifs, those predicted to form β strands result in lower mRNA levels than the remaining motifs (P < 0.05, Fig. 5C, right panel). To confirm the destabilizing role of the specific dipeptides identified in our study, we disrupted them by moving the bulky and positively charged amino acids to opposite ends without changing the amino acid composition in 1,079 endogenous motifs (Fig. 5D, top). As predicted, the resulting mutations increase mRNA levels (median log_2_ ΔmRNA = 0.38) with 783 mutated motifs having significantly higher mRNA levels (P < 0.05) than their wild-type counterparts (Fig. 5D, bottom). Examination of destabilizing motifs with annotated β strand structures in the Protein Data Bank (PDB) shows that these β strands are part of anti-parallel β sheets and are significantly longer than the 5-6 residue length of typical β strands (*42*) (Fig. 5E). Together, these results show that β-stranded endogenous motifs containing bulky and positively charged dipeptides can stall ribosomes and trigger mRNA instability.

## Discussion

Here, we identify a combinatorial code composed of bulky, positively charged, and extended β strand nascent peptides that regulates translation and mRNA stability in human cells. We demonstrate that a minimal combination of these sequence and structural elements is sufficient to induce ribosome stalling and cause changes in gene expression, and is widespread in the human proteome. As discussed below, elements of the code uncovered here allow us to synthesize a large body of observations on nascent peptide-mediated stalling of ribosomes and regulation of mRNA stability in human cells. Our results also point to a role for the ribosome as a post-synthesis filter against nascent peptide sequences that are bulky and aggregation prone.

While positive charge in the nascent peptide can slow ribosomes (*27, 28*), our results show that positive charge by itself is insufficient to induce changes in gene expression in human cells. The importance of bulky amino acids for mRNA stability effects is in line with the role of side chain bulk in ribosome-associated quality control in *S. cerevisiae* (*43*). Further, bulky synthetic amino acid analogs in the nascent peptide and small molecules that add bulk to the exit tunnel can both reduce ribosome elongation rate (*44*–*47*). Ribosome profiling in *S. cerevisiae* and human cells shows that tripeptide combinations of bulky and positively charged amino acids are enriched at sites of increased ribosome density (*23, 48*). Bulky and positively charged amino acids also play critical roles in many known ribosome-arresting peptides (*7, 49*–*52*). Structural studies of arrest peptides suggest that the bulky and positively charged amino acids might stall ribosomes by altering the geometry of the peptidyltransferase center (PTC) and/or by steric interactions with the constriction point in the exit tunnel formed by the uL4 and uL22 proteins (*7, 51, 53, 54*).

Our work shows that extended β strand motifs in nascent peptides contribute to ribosome stalling and mRNA instability in human cells. This role of a simple secondary structural motif like β strand is surprising given that cryo-EM studies of stalled ribosome nascent chain complexes reveal a diverse range of extended conformations, turns, and helices that are specific to each nascent arrest peptide (*7, 47, 52, 55, 56*). This comparison is complicated by the fact that cryo-EM studies are performed on post-arrest complexes where the nascent chain might have already undergone extensive conformational rearrangements. Further, while several destabilizing motifs uncovered here form β strands *in silico* and *in vitro* in isolation, they might have a significantly different structure within the confined geometry of the ribosome exit tunnel (*39, 57* –*59*). At the molecular level, β strands in nascent chains could contribute to stalling as an allosteric relay that communicates steric interactions between the nascent chain and the distal portions of the ribosome exit tunnel such as the uL4/uL22 constriction to the PTC (*45, 54, 55, 60*). This possibility is supported by our observation that destabilizing dipeptide repeats are at least 10-12 acids long (Fig. 2C), which is consistent with the distance between the uL4/uL22 constriction and the PTC.

The nascent peptide code for mRNA stability described here is significantly more complex and localized along the mRNA than previously associated factors such as codon, amino acid, and GC content (*12*–*15*). Thus, ribosome stalling nascent peptides might exert their effects on mRNA stability through distinct cellular pathways compared to the ones sensing codon, amino acid, and GC content of mRNAs (*22, 61*–*63*). Until our work, the UL4 CMV leader peptide, poly-lysine sequences encoded by poly-A, and the *Xbp1* arrest sequence were the only known nascent peptide motifs with ribosome stalling ability in human cells (*64*–*66*). However, the effect of these stall sequences on mRNA stability is not well characterized, with mild or no effects on steady state mRNA levels seen in transiently transfected reporter assays (*3, 67, 68*). Several other nascent peptide sequences stall human ribosomes in the presence of small molecule metabolites or drugs in the ribosome exit tunnel (*69, 70*). It is likely that the effect of nascent peptide motifs on ribosome stalling and mRNA stability is modulated by other co-translational events such as nascent protein folding outside the ribosome (*71, 72*), membrane insertion (*73, 74*), and multiprotein assembly (*75, 76*).

In addition to the sequence and structural determinants of nascent peptide-mediated ribosome stalling studied here, several classes of nascent peptide sequences that stall ribosomes might not be revealed by our assay. For example, poly-prolines do not emerge as destabilizing motifs in our assay even though they are known to stall ribosomes (*23*). This is likely because poly-proline stalls are resolved without triggering quality control or mRNA instability (*17*). While extended β strands are the primary structural motif associated with stalling here, we also find motifs with unstructured regions that are nevertheless destabilizing (Figs. 3A, 5B). This might be in part due to limitations of existing computational methods (*35*) to predict secondary structures or their limited relevance to secondary structures forming inside the ribosome. It is also likely that the combinatorial code of positively charged, bulky, and β strand sequences uncovered here underlies some, but not all, classes of nascent peptides that have the potential to stall ribosomes and effect changes in gene expression.

The nature of the nascent peptide code uncovered here has important implications for cellular homeostasis and disease. Ribosome stalling and mRNA destabilization induced by bulky and extended β strands, which are highly aggregation prone (*77, 78*), implies that the ribosome has an intrinsic ability to throttle the synthesis of such proteins. Such throttling might serve as a quality control mechanism to test the ability of β strands to fold into anti-parallel β sheets and thus avoid aggregation. This ribosomal selectivity filter acts before other cotranslational mechanisms such as codon optimality that help avoid aggregation after β strands emerge from the ribosome (*79, 80*). Finally, the gene regulatory potential of the destabilizing dipeptide motifs uncovered here suggests that disease-causing missense mutations occuring at these motifs might exert their phenotype by altering protein expression *in cis* rather than protein activity.

## Author Contributions

P.C.B. designed research, performed experiments, analyzed data, and wrote the manuscript. H.P. designed research and performed experiments. A.R.S. conceived the project, designed research, analyzed data, wrote the manuscript, supervised the project, and acquired funding.

## Acknowledgements

We thank members of the Subramaniam lab, the Zid lab, the Basic Sciences Division, and the Computational Biology program at Fred Hutch for discussions and feedback on the manuscript. This research was funded by NIH R35 GM119835, NSF MCB 1846521, and the Sidney Kimmel Scholarship received by ARS. This research was supported by the Genomics Shared Resource of the Fred Hutch/University of Washington Cancer Consortium (P30 CA015704) and Fred Hutch Scientific Computing (NIH grants S10-OD-020069 and S10-OD-028685). The funders had no role in study design, data collection and analysis, decision to publish, or preparation of the manuscript.

## Materials and Methods Plasmid construction

Plasmids, oligonucleotides, and cell lines used in this study are listed in Supplementary Tables S1-S3.

### Parent vector construction

The AAVS1-targeting parent vector pPBHS285 used for this study was constructed using Addgene plasmid #68375 (*81*) as a backbone. The PGK1 promoter was replaced with the CMV promoter and the native pCMV 5′ UTR region. The coding sequence was replaced by a codon-optimized mKate2 and eYFP fusion cassette, linked with two 2A linker sequences. These 2A sequences surround a cassette encoding an EcoRV restriction site, Illumina R1 sequencing primer binding site, and a T7 promoter. The R1 primer binding and and T7 sequences are in reverse orientation (3′ -5′) for *in vitro* transcription and sequencing of inserts and barcode sequences at the EcoRV site.

### Variable oligo pool design

Four oligo pools were designed for this study.

Pool 1 (Fig. 1B-E, Fig. 3A,C) encodes all possible dicodon (6 nt) combinations, for a total of 4096 codon pairs. These 6nt dicodon inserts were repeated eight times to create 8× dicodon repeat inserts, each 48nt in length.

Pool 2 (Fig. 2C,D, Fig. 4D) encodes several dipeptide combinations identified in Library 1 as causing mRNA instability. For Fig.2C, the number of dipeptide repeats was systematically reduced from 8 to 1. Repeats were replaced with a Ser-Gly linker, shown to be not destabilizing in Library 1, to maintain a constant 48nt insert length. For Fig. 2D, periodicity of dipeptides was by interspersing 1, 2, or 4 tandem repeats of each dipeptide with an equal number of its sequence-reversed counterpart. For Fig. 4D, destabilizing dipeptides KV and SF were combined and rearranged to form either alpha helices or beta strands, as predicted by S4PRED.

Pool 3 (Fig. 5C-E) encodes the 16 amino acid nascent peptide motifs from the human proteome identified as potentially destabilizing by the scoring method described in Fig.5A along with 4 flanking codons on either side. The library encodes the top 1079 predicted stalling motifs with a peptide score > 9, and 122 control motifs with a peptide score < 3. The library also includes the mutants with reordered amino acids from the 1,079 endogenous destabilizing dipeptide motifs, which were designed as described in Fig.5D.

Pool 4 (Fig 2A,B, Fig S3) encodes 8 inserts: 3 destabilizing dipeptide repeats (RH)_8_, (VK)_8_, (SF)_8_, their respective frameshifit controls (PS)_8_,(QS)_8_,(FQ)_8_, the β strand peptide (SVKF)_4_, and the α helix peptide (SKVF)_4_.

Oligo pools 1-3 were synthesized by Twist Biosciences with fanking sequences for PCR and cloning into the EcoRV site of the parent pPBHS285 vector. Oligo pool 4 was cloned by PCRing individual inserts and pooling them before cloning.

### Plasmid library construction

Parent vector pPBHS285 was digested with EcoRV. The oligo pools described above were PCR amplified using primers oHJ01 and either oPB348 (Library 1) or oPB409 (Libraries 2–4). oPB348 and oPB409 both encode a 24 nt random barcode region, comprised of 8×VNN repeats to exclude in-frame stop codons (where V is any nucleotide except T). Barcoded oligo pools were cloned into pPBHS285 by Gibson assembly. Assembled plasmid pools were transformed with high efficiency into NEB10Beta *E*.*coli*. For pools 1-3, the transformed plasmid pools were extracted from 15-50 *E*.*coli* colonies per insert in the library, thus bottlenecking the number of unique barcodes present in each plasmid pool. Resulting plasmid pools contained between 60,000–400,000 unique barcode sequences for pools 1-3. For pool 4, the transformed library was bottlenecked to around 150 barcodes per insert, and 6 such pools with distinct barcodes were extracted for multiplexed library preparation of different cell lines.

The plasmid libraries corresponding to pools 1-4 are pPBHS286, pPBHS309, pHPHS296, and pHPHS406, respectively. Variable insert and barcode sequences for each plasmid library are provided as part of the data analysis code.

### CRISPR vectors

The CLYBL-targeted Cas9-BFP expression vector pHPHS15 was constructed by Golden Gate assembly of either entry plasmids or PCR products with pHPHS11 (MTK0_047 (*82*) Addgene #123977) as backbone, pHPHS3 (MTK2_007 (*82*) Addgene #123702) for the pEF1a promoter, pADHS5 (*83*) (pU6-(BbsI)_CBh-Cas9-T2A-BFP (*84*) Addgene #64323) for the Cas9-2A-BFP insert cassette, and pHPHS6 (MTK4b_003 (*82*) Addgene #123842) for the rabbit β-globin terminator. sgRNA vectors pPBHS320 (gRNA_AAVS1-T1 Addgene #41817) and pADHS4 (*83*) (eSpCas9(1.1)_No_FLAG_AAVS1_T2 Addgene #79888) were used for insertion at the AAVS1 locus. pASHS16 (MTK234_030 spCas9-sgRNA1-hCLYBL (*82*) Addgene #123910) was used for insertion at the CLYBL locus.

### Cell line maintenance and generation

HEK293T cells (RRID:CVCL_0063, ATCC CRL-3216), HCT116 cells (RRID:CVCL_0291, NCI60 cancer line panel), and HeLa cells (RRID:CVCL_0030, ATCC CCL-2) were grown in DMEM (Thermo 11965084). K562 cells (RRID:CVCL_0004, ATCC CCL-243) were grown in IMDM (Thermo 12440053). Media for all cells was supplemented with 10% FBS (Thermo 26140079). Cells were grown at 37C in 5% CO2. All transfections into HEK293T, HCT116, and HeLa cells were performed using Lipofectamine 3000 (Thermo L3000015). Transfections into K562 cells were performed using an Amaxa Nucleofector V kit (Lonza VCA-1003). HEK293T cells that stably express Cas9 (hsPB80) were generated by transfecting the CLYBL::Cas9-BFP vector pHPHS15 and spCas9 sgRNA1 hCLYBL vector, and selecting with 200*μ*g/mL hygromycin.

### CRISPR integration of plasmid libraries

hsPB80 CLYBL::Cas9-BFP HEK293T cells were seeded to 50% confluency on 15 cm dishes for all library transfections. 10 *μ*g of library plasmid (pPBHS286, pPHBS309, or pHPHS296) and 1.5 *μ*g of each AAVS1 targeting CRISPR vector were transfected per 15 cm dish. pPBHS286, and pPBHS309 were each transfected into a single 15 cm dish. pHPHS296 was transfected into three 15 cm dishes. pHPHS406 pools with different barcodes were transfected into single 10 cm dishes of hsPB80, HeLa, HCT116 and 2 million cells of K562. Cells were selected with 2 *μ*g/mL puromycin, added 48 hours post-transfection. Cells from the three pHPHS296 transfections were combined at the start of selection. Puromycin selection was removed after 6-10 days, once cells were growing robustly in selection. 24 hours after removing puromycin selection, stable library cells were plated into two separate 15cm dishes, to reach 75% confluency the next day, for matched mRNA and gDNA harvests. For pHPHS406, libraries were maintained in two 10 cm dishes or T75 flasks (for K562).

### mRNA stability measurement

hsPB80 cells containing the stably integrated pHPHS406 library were seeded to 50% confluence in a 6-well plate. Actinomycin D (ActD) power was dissolved in DMSO at 1 mM (1.25 mg/mL) and added to each well of the 6-well plate to a final concentration of 5 *μ*g/mL. Before harvesting, 1 million HeLa cells containing the pHPHS406 library were lysed in 6 mL of Trizol reagent, to create a Trizol lysis solution containing a set number of mRNAs with different barcodes than those in the hsPB80 pHPHS406 pool, for barcode count normalization across samples. ActD treated hsPB80 wells were harvested at 0, 0.5, 1, 2, 4, and 6 hours after the addition of ActD by adding 0.75 mL of the Trizol lysis solution above to wells at each timepoint, then following the manufacturer’s Trizol mRNA extraction protocol.

### Library Genomic DNA extraction

Reporter library genomic DNA was harvested from one 75% confluent 15 cm or 10 cm dish of stably expressing library cells. Genomic DNA was harvested using Quick-DNA kit (Zymo D3024), following the manufacturer’s instructions, with 3 mL of genomic DNA lysis buffer per 15 cm plate, and 1 ml of the same buffer per 10 cm plate. Between 0.5-10 *μ*g of purified genomic DNA from each library sample was sheared into ∼350 nucleotide length fragments by sonication for 10 min on ice using a Diagenode Bioruptor. Sheared gDNA was then *in vitro* transcribed into RNA (denoted gRNA below and in analysis code) starting from the T7 promoter region in the insert cassette, similar to previous approaches (*85, 86*), using a HiScribe T7 High Yield RNA Synthesis Kit (NEB E2040S). Transcribed gRNA was treated with DNase I (NEB M0303S) and cleaned using an RNA Clean and Concentrator kit (Zymo R1013).

### Library mRNA extraction

Reporter library mRNA was harvested from one 75% confluent 15 cm or 10 cm dish of stably expressing library cells. mRNA was harvested by using 3 mL of Trizol reagent (Thermo) to lyse cells directly on the plate, and then following the manufacturer’s mRNA extraction protocol. Purified mRNA was then DNaseI (NEB M0303S) treated and cleaned using an RNA Clean and Concentrator kit (Zymo R1013).

### mRNA and genomic DNA barcode sequencing

Between 0.5-10 *μ*g of DNaseI-treated mRNA and gRNA for each library was reverse transcribed into cDNA using Maxima H Minus Reverse Transcriptase (Thermo EP0752) and a primer annealing to the Illumina R1 primer binding site (oPB354). A 170-nucleotide region surrounding the 24-nucleotide barcode was PCR amplified from the resulting cDNA in two rounds, using Phusion Flash High-Fidelity PCR Master Mix mastermix (Thermo F548L). Round 1 PCR was carried out for 10 cycles, with cDNA template comprising 1/10th of the PCR reaction volume, using primers oPB361 and oPB354. Round 1 PCRs were cleaned using a 2× volume of Agencourt Ampure XP beads (Beckman Coulter A63880) to remove primers. Cleaned samples were then used as template for Round 2 PCR, carried out for 5-15 cycles, using a common reverse primer (oAS111) and indexed forward primers for pooled high-throughput sequencing of different samples (oAS112-135 and oHP281-290). Amplified samples were run on a 1.5% agarose gel and fragments of the correct size were purified using ADB Agarose Dissolving Buffer (Zymo D4001-1-100) and UPrep Micro Spin Columns (Gene-see Scientific 88-343). Concentrations of gel-purified samples were measured using a Qubit dsDNA HS Assay Kit (Q32851) with a Qubit 4 Fluorometer. Samples were sequenced using an Illumina HiSeq 2500 or Illumina NextSeq 2000 in 1×50, 2×50, or 1×100 mode (depending on other samples pooled with the sequencing library).

### Insert-barcode linkage sequencing

Plasmid library pools 1-4 (pPBHS286, pPBHS309, pHPHS296, and pHPHS406) were diluted to 10 ng/*μ*L. A 240-nucleotide region surrounding the 48-nucleotide variable insert sequence and the 24-nucleotide barcode was PCR amplified from these pools in two rounds, using Phusion Flash High-Fidelity PCR Master Mix mastermix (Thermo F548L). Round 1 PCR was carried out for 10 cycles, with 10 ng/*μ*L plasmid pool template comprising 1/10th of the PCR reaction volume, using primers oPB29 and oPB354. Round 1 PCRs were digested with DpnI (Thermo FD1704) at 37°C for 30 minutes to remove template plasmid and cleaned using a 2× volume of Agencourt Ampure XP beads (Beckman Coulter A63880) to remove primers and enzyme. Cleaned samples were used as template for Round 2 PCR, for 5 cycles, using oAS111 and indexed forward primers (oAS112-135 and oHP281-290). Amplified Round 2 PCR products were purified after size selection and quantified as described above for barcode sequencing. Samples were sequenced using an Illumina MiSeq or Illumina NextSeq 2000 in 2×50 or 1×100 mode.

### Rabbit reticulocyte nanoluciferase transit time assay

DNA fragments encoding 4×KVFS and 4×SKVF (α helix) and 4×VKFS and 4×SVKF (β strand) peptides were generated by PCR-amplifying overlapping oligos that encode each sequence in the forward and reverse direction (oPB470-473 and oPB488-491). Nanoluciferase cassette was amplified from an IDT gBlock (oPN204) using oAS1287 and oPB465. Insert sequences and the Nanoluciferase cassette were combined by overlap PCR using oPB464 and oPB462, which add a 5′ T7 promoter site and a 3′ polyA tail to the amplified reporter template, with oAS1545 used to bridge oPB462 annealing. Resulting insert-Nanoluciferase cassette sequences were confirmed by Sanger sequencing. The PCR products were transcribed into mRNA using a HiScribe T7 High Yield RNA Synthesis Kit (NEB E2040S). mRNA was cleaned using an RNA Clean and Concentrator kit (Zymo R1013). *In vitro* Nanoluciferase reporter translation reactions were performed as described in Susorov et al. 2020 (*87*). Reaction mixture containing 50% of nuclease-treated rabbit reticulocyte lysate (RRL) (PRL4960, Promega) was supplemented with 30 mM Hepes-KOH (pH = 7.5), 50 mM KOAc, 1.0 mM Mg(OAc)_2_, 0.2 mM ATP and GTP, 0.04 mM of 20 amino acids (PRL4960, Promega), and 2 mM DTT. Nanoluciferase substrate furimazine (PRN1620, Promega) was added to the mixture at 1%. 15 *μ*L aliquots of the mixture were placed in a 384-well plate and incubated at 30°C for 5 min in a microplate reader (Tecan INFINITE M1000 PRO). Translation reactions were started by simultaneous addition of 3 *μ*L mRNA, to a final concentration of 10 ng/*μ*L, and luminescence signal was recorded every 10 seconds over a period of 25 minutes.

### Circular dichroism

4×SKVF (α helix) and 4×SVKF (β strand) peptides were commercially synthesized (Genscript) at >90% purity level. Peptides were dissolved in water to 400 *μ*M concentration, then diluted into 10 mM sodium-phosphate buffer (pH = 7.5) and 0, 20, or 40 volumetric percent of 2,2,2-trifluoroethanol (TFE) to final concentrations ranging between 15-30 *μ*M. CD spectra were measured at 25C using a Jasco J-815 Circular Dichroism Spectropolarimeter. The CD spectra were recorded between 180-260 nm with a resolution of 0.5 nm for both peptides and blank buffer solutions in 1 mm cuvettes.

### Computational analyses

Pre-processing steps for high-throughput sequencing were implemented as `Snakemake` workflows (*88*). `Python` (v3.7.4) and `R` (v3.6.2) programming languages were used for all analyses unless mentioned otherwise.

In the description below, files ending in `.py` refer to `Python` scripts and files ending in `.Rmd` or `.R` refer to `R Markdown` or `R` scripts.

### Barcode to insert assignment

The raw data from insert-barcode linkage sequencing are in `.fastq` format. If the inserts and barcodes were on paired-end reads instead of singleend reads, the reads were renamed in increasing numerical order starting at 0 to enable easy matching of insert and barcode reads. This was done in `rename_fastq_paired_reads.py`. The oligo pools were used to create a reference `fasta` file in `create_reference_for_aligning_library.R`. A `bowtie2` (*89*) (v2.4.2) reference was created from the `fasta` file using the `bowtie2-build` command with default options. The insert read was aligned to the `bowtie2` reference using `bowtie2` command with options `-N 1 -L 22 --end-to-end` with the `--trim5` and `--trim3` options set to include only the region corresponding to the insert. The alignments were sorted and indexed using `samtools` (*90*) (v1.11) commands `sort` and `index` with default options. The alignments were filtered to include only reads with simple `cigar` strings and a `MAPQ` score greater than 20 in `filter_alignments.R`. The barcodes corresponding to each filtered alignment were parsed and tallied in `count_barcode_insert_pairs.py`. Depending on the sequencing depth, only barcodes that were observed at least 4-10 times were included in the tally. The tallied barcodes were aligned against themselves using `bowtie2-build` with default options and `bowtie2` with options `-L 24 -N 1 --all --norc`. The self-alignment was used to exclude barcodes that are linked to distinct inserts or ones that are linked to the same barcode but are aligned against each other by `bowtie2`. In the latter case, the barcode with the lower count is discarded. The final list of insert barcode pairs is written as a tab-delimited `.tsv.gz` file for aligning barcodes from genomic DNA and mRNA sequencing below.

### Barcode counting in genomic DNA and mRNA

The raw data from sequencing barcodes in genomic DNA and mRNA is in `.fastq` format. The filtered barcodes `.tsv.gz` file from the insert-barcode linkage sequencing is used to create a reference `fasta` file in `create_bowtie_reference.R`. A `bowtie2` (v2.4.2) reference was created from the `fasta` file using the `bowtie2-build` command with default options. The barcodes were aligned to the `bowtie2` reference using `bowtie2` command with options `-N 1 -L 20 --norc` with the `--trim5` and `--trim3` options set to include only the region corresponding to the barcode. The alignments were sorted, indexed, and tallied using the `samtools` commands `sort`, `index`, `idxstats` with default options. GNU `awk` (v4.1.4) was used for miscellaneous processing of tab-delimited data between preprocessing steps. The final list of counts per barcode in each sample of genomic DNA or mRNA is written as a tab-delimited `.tsv.gz` file for calculating mRNA levels below.

### mRNA quantiication

All barcode counts corresponding to each insert in each sample were summed. Only inserts with a minimum of 200 reads and 6 barcodes summed across the mRNA and gRNA samples were included. Otherwise the data were designated as missing. mRNA levels were calculated as the log2 ratio of the summed mRNA barcode counts to the summed gRNA barcode counts. mRNA levels were median-normalized within each library. For mRNA stability measurements, the summed mRNA counts for each insert at each time point were normalized by the total barcode counts for the spiked-in HeLa cells at the same time point. Then, the spike-in normalized mRNA levels for each insert were further normalized to the time 0 value.

### Linear statistical modeling of mRNA levels

Amino acid scales for isoelectric point pI, bulkiness, and secondary structure propensity were taken from prior studies (*32, 36, 91*). The median-normalized mRNA levels for lysine, arginine, or glutamate dipeptides were modeled as a function of amino acid scales (as indicated in the figures) using the `R` function `lm` with default parameters. Only fit coefficients significantly different from zero (P < 0.05) are reported for each linear model.

### Secondary structure prediction

Secondary structure was predicted solely from the amino acid sequence using the default single sequence model in S4PRED (*35*) (downloaded from https://github.com/psipred/s4pred on Apr 17, 2021) and the neural network was used without any modification in `predict_secondary_structure.py`.

Cartoons of 4×SVKF and 4×SKVF in Fig. 4A were predicted using the PEP-FOLD3 server (*92*) with default parameters and the resulting PDB files were visualized using PyMOL software (Schrodinger).

### Calculation of secondary structure content from circular dichroism

The raw circular dichroism data (Fig. 4B, left panel) were converted to the two-column spectrum file format as required for `SESCA` (*93*) (v095, downloaded from https://www.mpibpc.mpg.de/sesca on Jul 28, 2021). Secondary structure was estimated using the `SESCA` script `SESCA_deconv.py` using the pre-computed basis set `Map_BB_DS-dTSC3.dat` and options @err 2 @rep `100`. The output `.txt` file was parsed to extract the α helix, β strand, and random coil content shown in Fig. 4B, right panel.

### Calculation of ribosome transit time

The raw luminescence vs. time data (Fig. 4C, middle panel) were fit to a straight line in the linear regimes (600s < t < 900s for 4×SKVF and 4×KVFS, 900s < t < 1200s for 4×SVKF and 4×VKFS) using the `R` function `lm`. The `intercept` term from the fit was used as the transit time of ribosomes across the full transcript and its mean and standard error across technical replicates is shown in Fig. 4C, right panel.

### Calculation of ribosome density around endogenous destabilizing motifs

One canonical transcript for each protein coding gene from `GENCODE v32` was identified by filtering for transcripts that are annotated as `CCDS` or that have a `transcript_support_level` of 1. Duplicate transcripts on the Y chromosome were discarded. After the above filtering, the transcript with the smallest ENSEMBL transcript id was picked for each gene, followed by the transcript with the longest CDS, and finally by the transcript with the smallest CCDS id. The same procedure was repeated for identifying canonical mouse transcripts from `GENCODE vM25`. The translated CDS of canonical transcripts were scanned for dipeptides formed by all 20 combinations of `K`, `R` and `L`, `I`, `Y`, `V`, `F`. The location of such dipeptides was given a score of 1, and locations of overlapping dipeptides were given a score of 2, and a score of 0 otherwise. The center of the 16 amino acid window with the highest peptide score was identified using the `convolve` function from the `Python numpy` package with the option `mode` set to `same` by convolution with a vector of length 16 filled with 1, followed by the `argmax` function to find the location of the maximum convolution score. A control motif in each transcript was randomly identified among the subset of 16 amino windows that had a peptide score less than 3. The above identification of destabilizing and control motifs is done in `find_destabilzing_motifs.py`.

Ribo-seq data from the indicated studies (*23, 40, 41*) were downloaded from SRA based on GEO accession numbers (GSM39075{97,98}, GSM28281{65,66,67}, GSM16440{76,77,78,79,88,89}), trimmed to remove 5′ and 3′ adapters using `cutadapt` (*94*), filtered to remove contaminating ribosome RNA reads, aligned against the above canonical transcripts using `bowtie2` with default parameters, sorted and indexed using `samtools`. The alignments were trimmed to the P-site as specified in the studies and tallied to calculate coverage at each transcript location (`calculate_transcripts_coverage.R`). The list of transcripts was filtered for transcripts that had an average coverage of 1 read per codon in the CDS region. The P-site density around each motif was calculated by normalizing the read counts at each location by the average read density on that transcript (`find_ribosome_density_around_motifs.py`). The normalized P-site densities were averaged at each location around the motif across all destabilizing or control motifs to obtain the metagene profiles shown in Figs. 5B, S4A, S4B (`plot_destabilizing_motif_density.Rmd`).

### Statistical analyses

For barcode sequencing, error bars were calculated as the standard deviation of 100 bootstrap samples of barcodes across the gRNA and mRNA samples. The standard deviation was measured for the log2 mRNA levels calculated as described in the *mRNA quantification* section. For all other experiments, the standard error of the mean was calculated using the `std.error` function from the `plotrix R` package. P-values for statistical significant differences were calculated using the `t.test` or `wilcox.test R` functions as appropriate for each figure (see figure captions).

**Figure S1.**
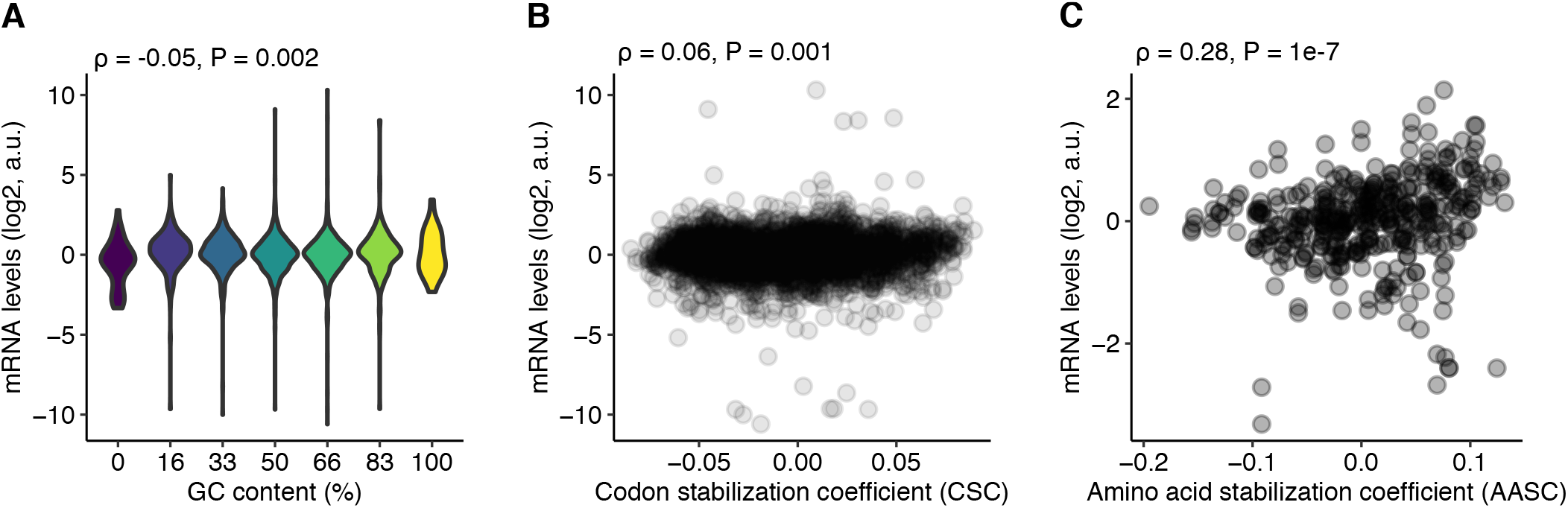
**(A)** mRNA levels of dicodon repeats as a function of their GC content. **(B)** mRNA levels of dicodon repeats as a function of their mean codon stabilization coefficient (*12*). **(C)** mRNA levels of dipeptide repeats as a function of their mean amino acid stabilization coefficient (*15*). Spearman correlation coefficient ρ and its P-value are shown in A-C.

**Figure S2.**
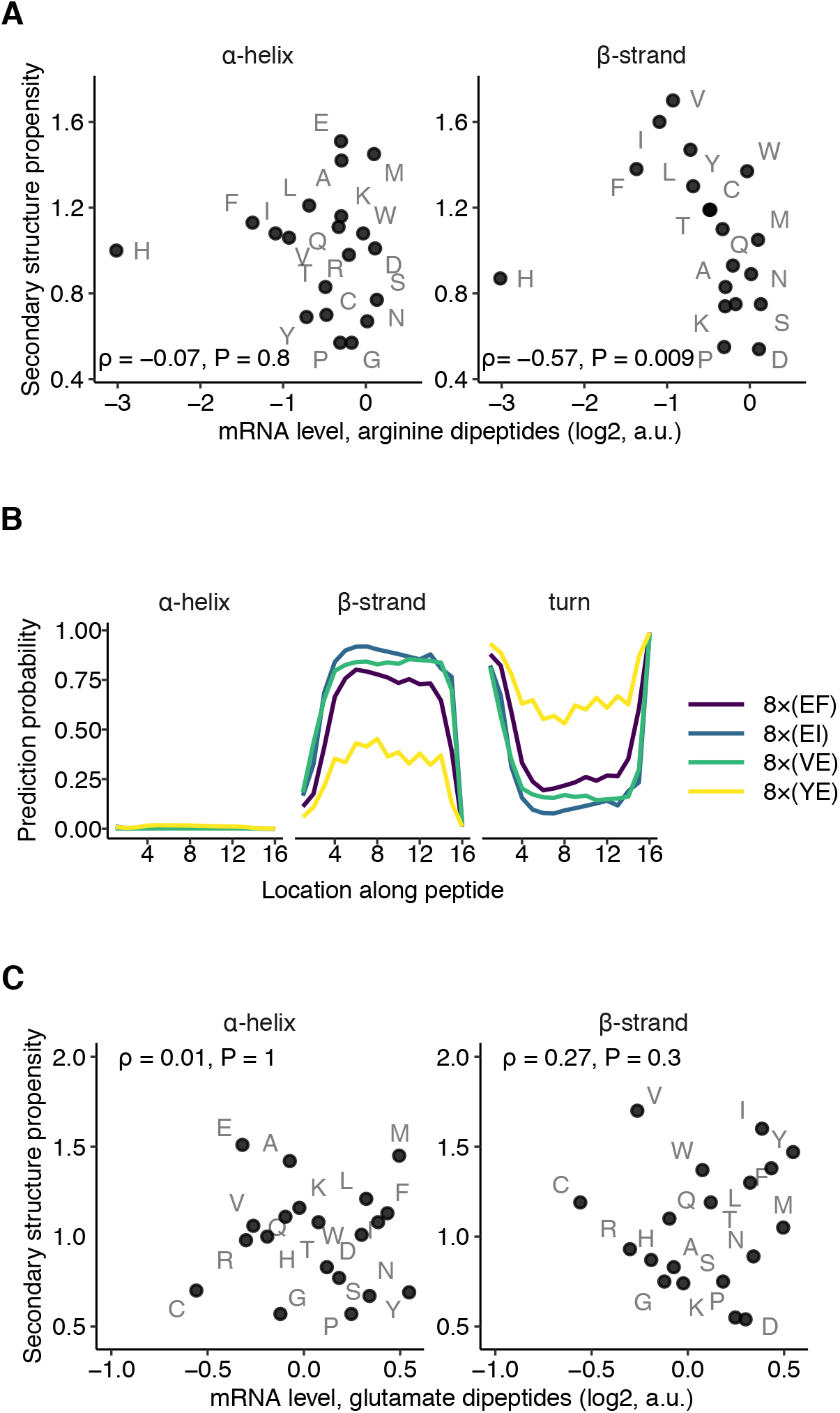
(**A**,**C)** mRNA levels of dipeptide repeat-encoding reporters with arginine (A) or glutamate (C) in one position and one of twenty amino acids in the other position of the repeat (labeled in grey) plotted as a function of the propensity (*36*) of the second amino acid to occur in a β strand (right) or an α helix (left). ρ is the Spearman correlation coefficient between the two axes with the indicated *P* value. **(B)** Computationally predicted secondary structure probability along 16 amino acid-long peptide sequences encoded by dipeptides with glutamate that form β strands. Secondary structure probabilities are predicted using S4PRED (*35*). Amino acids are labeled by their one-letter codes.

**Figure S3.**
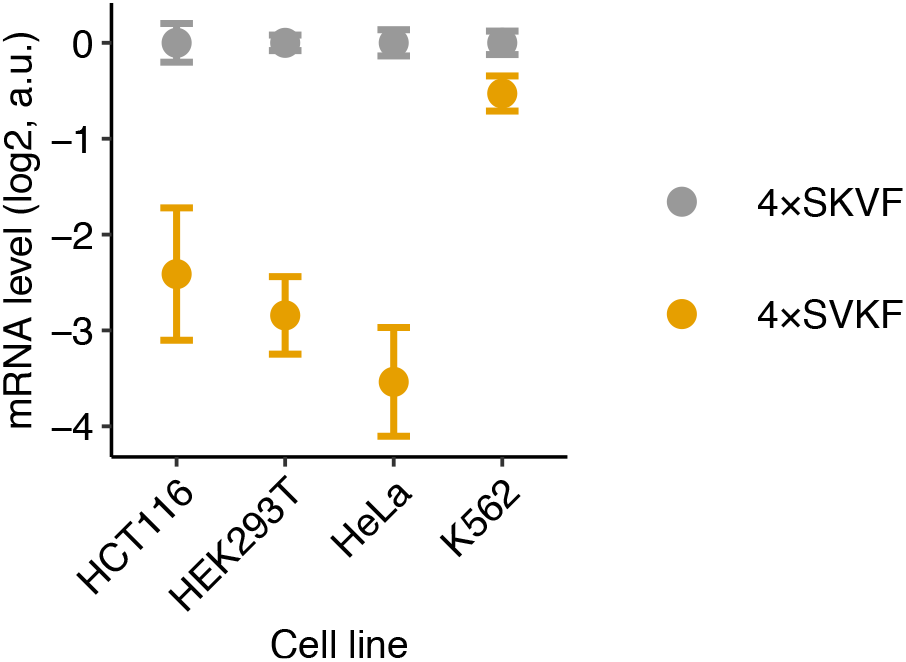
mRNA levels of reporters with α helix (SKVF)_4_ or β strand (SVKF)_4_ forming peptides in 4 different cell lines HCT116, HEK293T, HeLa, and K562. Error bars represent standard errors of measurement over barcoded replicates calculated by bootstrap.

**Figure S4.**
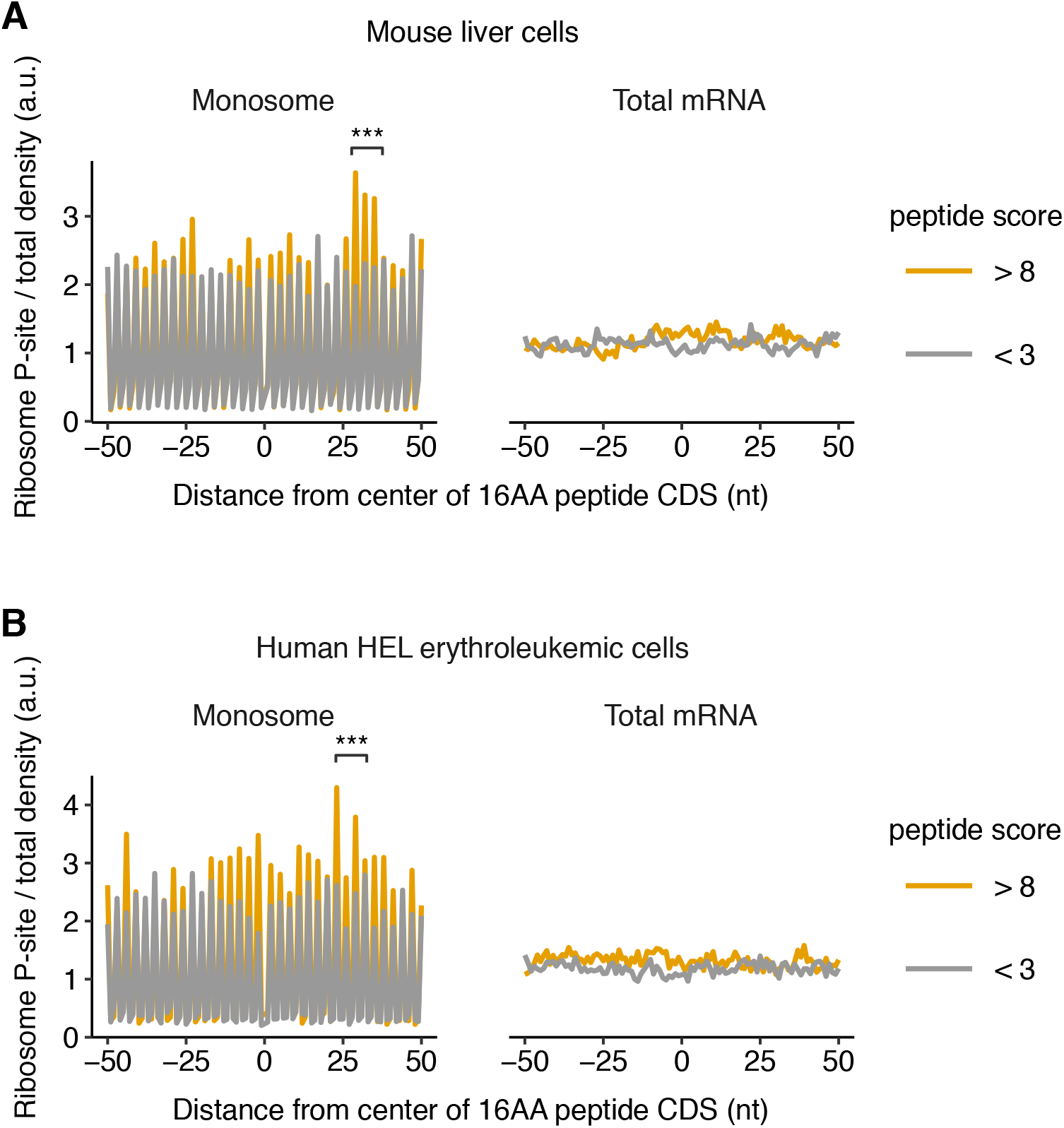
**(A**,**B)** Measured ribosome P-site density or total RNA density around destabilizing dipeptide motifs in mouse liver cells (*41*) and human HEL erythroleukemic cells (*40*). The first nucleotide of the 9th codon encoding the 16AA peptide motif is designated as 0nt. ^***^: P < 0.001 (two-sided Mann-Whitney U test) for increase in ribosome P-site density in +25 to +35 (A) or +20 to +30nt (B) around the destabilizing peptide motif. Comparison is between motifs with peptide score > 8 and randomly chosen control motifs in the same set of genes with peptide score < 3.

